# The C-terminus of α-Synuclein regulates its dynamic cellular internalization by Neurexin 1β

**DOI:** 10.1101/2023.07.28.550969

**Authors:** Melissa Birol, Isabella Ioana Douzoglou Muñoz, Elizabeth Rhoades

## Abstract

The aggregation of the disordered neuronal protein, α-Synuclein (αS), is the primary pathological feature of Parkinson’s disease. Current hypotheses favor cell-to-cell spread of αS species as underlying disease progression, driving interest in identifying the molecular and cellular species involved in cellular internalization of αS. Prior work from our lab identified the chemically specific interaction between αS and the pre-synaptic adhesion protein neurexin 1β (N1β) to be capable of driving cellular internalization of both monomer and aggregated forms of αS. Here we explore the physical basis of N1β-driven internalization of αS. Specifically, we show that spontaneous internalization of αS by SH-SY5Y and HEK293 cells expressing N1β requires essentially all of the membrane-binding domain of αS; αS constructs truncated beyond residue 90 bind to N1β in the plasma membrane of HEK cells, but are not internalized. Interestingly, prior to internalization, αS and N1β co-diffuse rapidly in the plasma membrane. αS constructs that are not internalized show very slow mobility themselves, as well as slow N1β diffusion. Finally, we find that truncated αS is capable of blocking internalization of full-length αS. Our results draw attention to the potential therapeutic value of blocking αS-N1β interactions.

## Introduction

Alpha-synuclein (αS) is a small (140 residues), intrinsically disordered neuronal protein (Davidson et al., 1998; Theillet et al., 2016). In Parkinson’s disease, αS is the primary component of intracellular fibrillar aggregates that are the hallmark of disease (Galvin et al., 1999; Spillantini et al., 1998). Natively, αS is primarily localized to pre-synaptic termini (Kahle et al., 2000; Maroteaux et al., 1988), where it binds reversibly to cellular membranes (Bendor et al., 2013; Bussell and Eliezer, 2003; Fortin et al., 2004; Kahle et al., 2000). Because of its central role in Parkinson’s disease and other synucleinopathies, αS is of major biomedical interest.

αS has three domains (Figure 1): (1) the first ∼90 residues consist of seven, 11 residue imperfect repeats of a KTKEGV motif which mediate membrane binding through formation of a long, amphipathic helix (Bussell and Eliezer, 2003; Ferreon et al., 2009; Jao et al., 2004; Trexler and Rhoades, 2009); all 6 (A30P, E46K, H50Q, G51D, A53T/E) mutations linked to familial forms of Parkinson’s disease are found in this region; (2) the central non-amyloid component (NAC) domain between residues 61-95 which also makes up the core of aggregated synuclein fibers (Guerrero-Ferreira et al., 2018; Li et al., 2018; Tuttle et al., 2016) is encompassed within the membrane binding domain; and (3) the highly acidic C-terminus. Like many intrinsically disordered proteins, αS is subject to numerous post-translational modifications, several of which are of interest as biomarkers or for putative roles in αS aggregation in disease (Beyer, 2006; Beyer and Ariza, 2013; Kellie et al., 2014). While the majority of identified modifications to αS are found on only a fraction of the total protein (Kellie et al., 2014), N-terminal acetylation has been identified on virtually all αS derived from human or other mammalian cells and tissue (Anderson et al., 2006) and thus is a more physiologically relevant form of the protein than αS lacking this modification.

**Figure 1:**
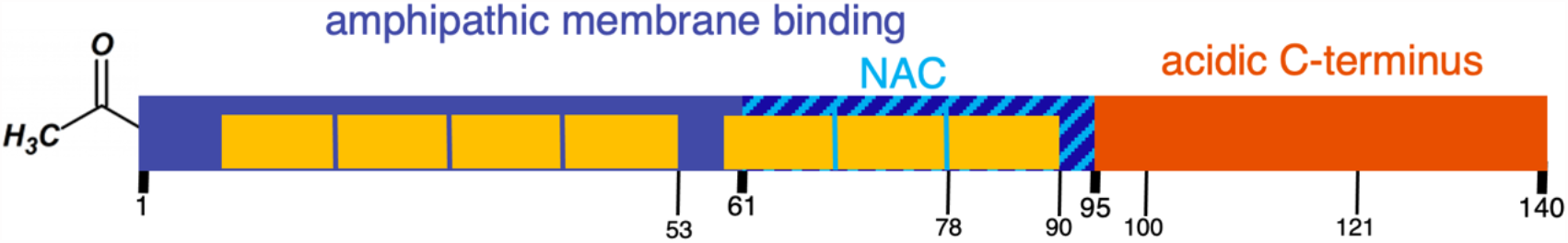
Schematic of αS_acetyl_. Three major domains are labeled on the schematic and the seven, 11 residue membrane binding repeats are shown in yellow. Also marked are the truncations created for this study. For all constructs, a S9C mutation was made to allow for site-specific labeling.

While αS is primarily cytoplasmic, it is also found extracellularly and in cerebral spinal fluid (Bieri et al., 2018; Kang et al., 2016; Lee et al., 2005; van Steenoven et al., 2018; Zambon et al., 2019). In Parkinson’s disease, it has been proposed that propagation of pathological aggregates occurs via a prion-like mechanism, where toxic αS assemblies are released into the extracellular matrix by ‘infected’ neurons (Guo and Lee, 2014). These species are internalized by neighboring neurons, seeding aggregation of the endogenous αS in these cells. Although it lacks a canonical secretion sequence, cell culture models show that monomeric αS is continuously secreted to the extracellular media (Emmanouilidou et al., 2010; Yamada and Iwatsubo, 2018) where it may be re-internalized. Consequently, there is significant interest in identifying and characterizing the molecular components and cellular pathways involved in the internalization of αS. A number of plasma membrane proteins have been identified as αS receptors (Birol et al., 2019; Diaz-Ortiz et al., 2022; Mao et al., 2016; Shrivastava et al., 2015; Urrea et al., 2017). Prior work from our lab demonstrated that expression of the neuronal adhesion protein, neurexin-1β (N1β), in HEK293 cells is sufficient to cause spontaneous internalization of both monomer and fibrillar αS (Birol et al., 2019). Moreover, this uptake showed chemical selectivity in that it was dependent both upon the N-terminal acetylation of αS as well as the N-linked glycan on the globular extracellular domain of N1β (Birol et al., 2019).

As noted above, various neuronal cell surface receptors have been identified for different forms of αS – monomer, oligomer and fibrillar - pointing to the existence of multiple internalization pathways. There is also evidence that the inhibition of a single cellular endocytosis pathway is not sufficient to completely stop cellular entry of αS. Our current understanding of the molecular elements governing the internalization of αS is limited. Here we investigate the molecular features associated with N1β-driven internalization of αS_acetyl_. Specifically, we used truncated variants of αS_acetyl_ to identify regions involved in N1β-mediated binding and uptake in HEK 293 cells. We show that full-length and truncated αS_acetyl_ bind equally to HEK-N1β; however, they are not internalized to a similar degree. Specifically, we identify that the entire membrane-binding domain of αS_acetyl_ is essential for N1β-mediated internalization of αS_acetyl_. Moreover, we measured the mobility of αS_acetyl_/N1β complexes in the plasma membrane, prior to internalization, and found a correlation between the degree of mobility and the extent of uptake. Lastly, we report on a specific αS truncation that inhibits N1β-mediated internalization of full-length protein, opening a new avenue for therapeutic intervention focused on limiting the extracellular propagation of αS_acetyl_.

## Results & Discussion

### Robust spontaneous internalization of αS_acetyl_ requires the entire membrane-binding domain

Full-length N-terminally acetylated αS (^1-140^αS_acetyl_) was obtained by co-expression of αS with the NatB complex from saccharomyces cerevisiae in e coli (Birol et al., 2019; Johnson et al., 2010; Trexler and Rhoades, 2012). Truncated constructs of αS were created by introduction of a stop codon at the appropriate location in the plasmid encoding full-length αS. ^1-121^αS_acetyl_ and ^1-100^αS_acetyl_ were chosen because truncations near these residues have been identified in human tissue (Anderson et al., 2006; Bhattacharjee et al., 2019; Kellie et al., 2014); ^1-90^αS_acetyl_, ^1-78^αS_acetyl_ and ^1-53^αS_acetyl_ were selected because they correspond to fragments spanning 7, 6 and 4 of the membrane binding repeats, respectively (Figure 1). A serine to cysteine mutation was made at residue 9 to allow for site-specific labeling with Alexa 594 (AL594) maleimide. We have fluorescently labeled αS at this position with several different fluorophores and found minimal impact on its properties in a number of studies (Middleton and Rhoades, 2010; Nath et al., 2012; Sevcsik et al., 2011; Trexler and Rhoades, 2009; 2010).

SH-SY5Y cells are a neuroblastoma-derived immortal line that are frequently used as a model system for studies involving αS because they maintain many of the pathways that are dysregulated in Parkinson’s disease (Krishna et al., 2014). Moreover, they spontaneously internalize both monomer and pre-formed fibrillar (PFF) αS (Birol et al., 2019; Rodriguez et al., 2018). SH-SY5Y cells were incubated with 200 nM AL594 αS_acetyl_. Following 12 hours incubation, the media was exchanged to remove any remaining αS_acetyl_ and the cells were imaged. Internalized αS_acetyl_ can be seen as punctate structures in the cell body (Figure 2A), which we know from our prior work to be acidic endosomes or lysosomes (Birol et al., 2019). Comparable levels of uptake were found in cells incubated for 12 hours with ^1-140^αS_acetyl_, ^1-121^αS_acetyl_ and ^1-100^αS_acetyl_, while cells incubated with ^1-78^αS_acetyl_ and ^1-53^αS_acetyl_ did not show significant internalization (Figure 2A, B).

**Figure 2:**
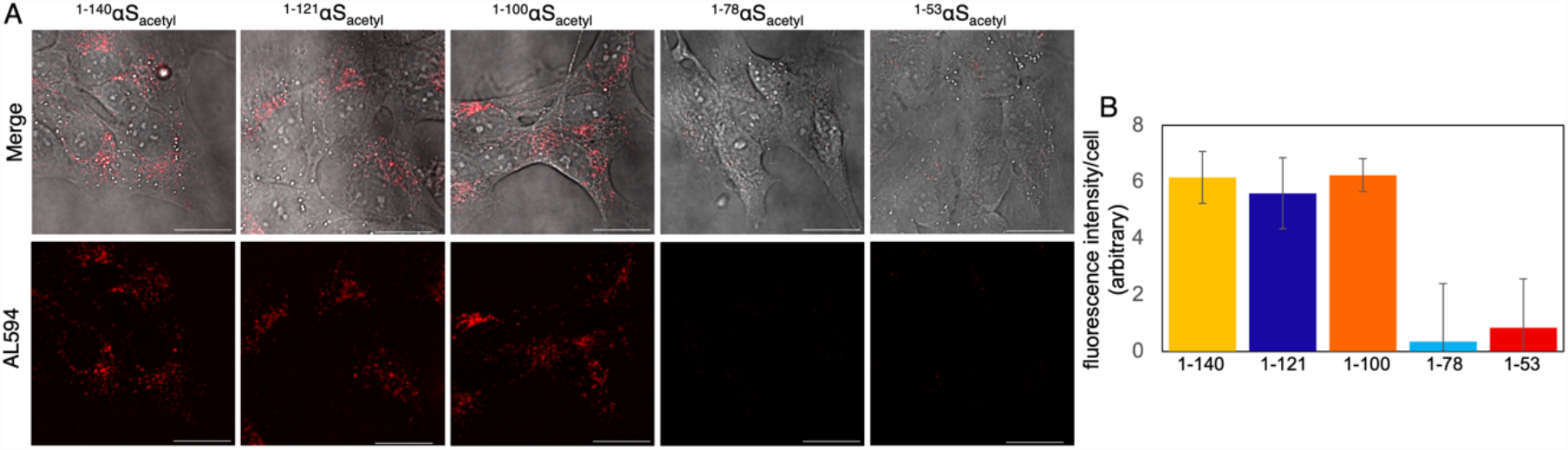
Internalization in SH-SY5Y is reduced for truncated αS_acetyl_ constructs. (A) Representative images of SH-SY5Y cells following 12-hour incubation with AL594 αS_acetyl_ variants. The fluorescence images are shown with (upper) and without (lower) DIC merge. (B) Quantification of extent of spontaneous internalization of αS_acetyl_ constructs by SH-SY5Y cells. Images obtained after 12-hour incubation with protein and quantified by total fluorescence intensity per cell. Analysis based on n= 100 cells, 3 independent experiments. Scale bars = 20 μm.

### Truncated αS_acetyl_ binds HEK-N1β

In contrast to SH-SY5Y cells, HEK293 cells do not spontaneously internalize ^1-140^αS_acetyl_ (Birol et al., 2019). Expression of N1β in HEK293 cells, however, is sufficient to drive αS_acetyl_ uptake (Birol et al., 2019). N1β with a C-terminal tGFP tag (N1β-tGFP) was expressed in HEK293 cells (HEK-N1β), where it localizes to the plasma membrane (Figure S1), and incubated 200nM AL594 ^1-140^αS_acetyl_ for various amounts of time. Following 1 h incubation, both N1β and ^1-140^αS_acetyl_ can be visualized primarily on the plasma membrane (Figure 3A). By 6 h, there is significant internalization of both proteins, where they co-localize in intracellular puncta (Figure 3C). Co-localization of the intracellular proteins is observed at 12 h and 24 h, as well, although there is some slight separation of N1β and αS_acetyl_ at the 24 h timepoint suggesting that their processing pathways may be diverging (Figure 3A, C). We observe a significant increase in the fraction of N1β signal on the cell membrane at 24 hours (Supplemental Figure S2), while ^1-140^αS_acetyl_ remain as intracellular punctate structures (Figure 3A, B); this may be the result of internalized N1β being trafficked back to the plasma membrane, or newly expressed N1β reaching the plasma membrane. Correlation analysis of these images confirms a decrease in the αS_acetyl_/ N1β co-localization signal at 24 h (Figure 3 Table).

**Figure 3:**
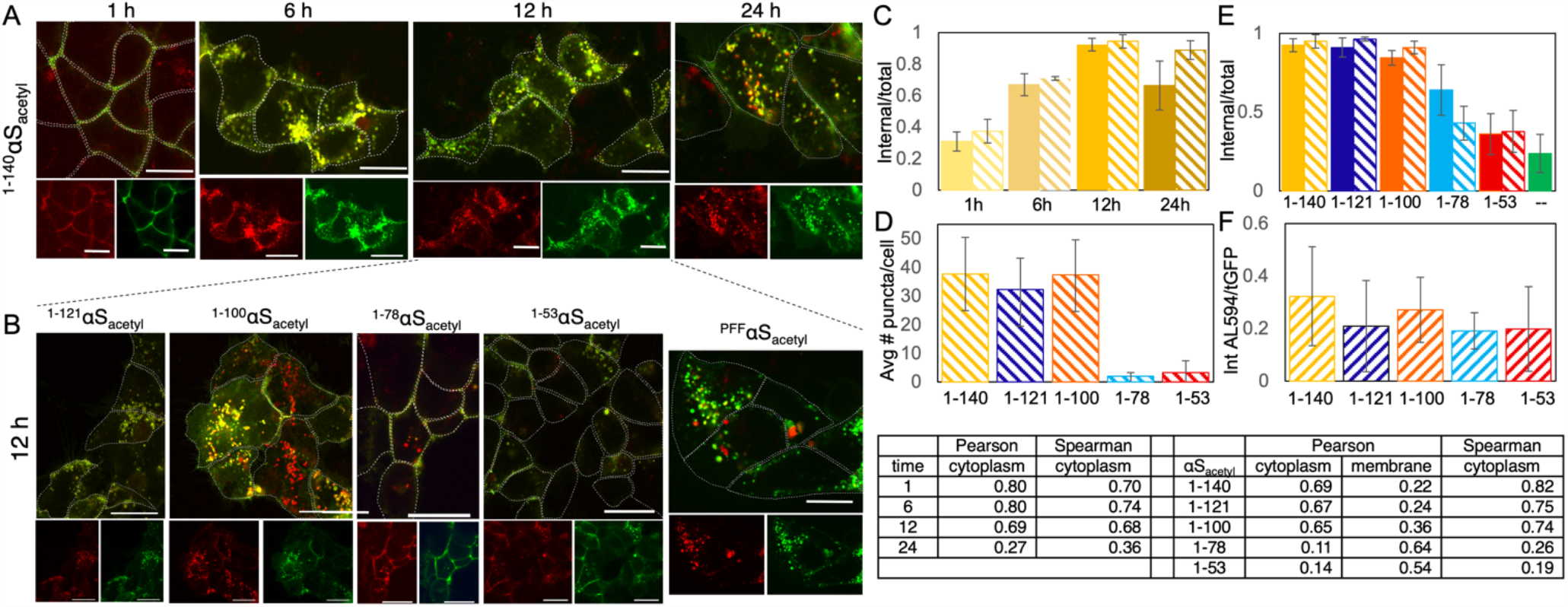
Internalization in HEK-N1β is reduced for truncated αS_acetyl_ constructs. (A) Representative images of HEK-N1β-tGFP cells following 1h, 6h, 12h, and 24h incubation with AL594 ^1-140^αS_acetyl_. The αS (red) and N1β (green) channels are shown separately below a larger image of their merge; cells are outlined in white dashed lines. (B) Representative images of HEK-N1β-tGFP cells following 12h incubation with AL594 αS_acetyl_ truncations or ^1-140^αS_acetyl_ PFFs. The αS (red) and N1β (green) channels are shown separately below a larger image of their merge; cells are outlined in white dashed lines. (C) Internalization quantification for (A) of N1β−tGFP (solid) and AL594 ^1-140^αS_acetyl_ (stripes) at the indicated incubation time. Colocalization was analyzed with the Pearson or Spearman correlation coefficient (Table). A larger coefficient reflects more overlap between the AL594 αS_acetyl_ and N1β−tGFP signals. Correlation coefficients were computed using the ImageJ plugin for colocalization. (D) Quantification of internalization for (B) of AL594 αS_acetyl_ truncations by HEK-N1β cells following 12 h incubation by puncta analysis. (E) Fraction of fluorescence signal of N1β-tGFP (solid) or AL594 αS_acetyl_ (stripes) from internalized proteins following 12 hours incubation with AL594 αS_acetyl_ or control (--) (F) Quantification of binding of αS_acetyl_ constructs to HEK-N1β-tGFP treated with an endocytosis inhibitor to prevent internalization. Table - Colocalization analysis of data from (C) and (E). Analysis for internalization and binding based on n = 100 cells, 3 independent experiments. Scale bars = 20 μm. Colocalization analysis based on n = 25 – 50 cells, 3 independent experiments.

At 12 h incubation, internalized αS_acetyl_ and N1β-tGFP co-localize with LysoTracker Deep Red, which labels lysosomes and acidified endosomes (Supplemental Figure S3). In experiments with the addition of control buffer lacking αS_acetyl_, the fraction of the cytoplasmic N1β is ∼0.25 at 12 hours (Figure 3E), comparable to what is observed in a 1 hour control (Supplemental Figure S2), indicating that there is no time-dependent behavior of the N1β signal in the absence of αS_acetyl_ treatment. We also created a construct of N1β bearing an N-terminal tGFP tag, tGFP-N1β; this tag precedes the signal sequence of N1β? and, as expected, when expressed in HEK293 cells, can be visualized throughout the cell body rather than localized to the plasma membrane (Supplemental Figure S1). HEK cells expressing tGFP-N1β do not spontaneously internalize ^1-140^αS_acetyl_ (Supplemental Figure S1), further underscoring the importance of the plasma membrane localization of N1β in driving uptake.

For comparison amongst the truncated αS_acetyl_ constructs, the 12 h incubation timepoint was chosen. As with the SH-SY5Y cells (Figure 2), when quantified by the fluorescence of intracellular puncta, internalization of ^1-121^αS_acetyl_ and ^1-100^αS_acetyl_ was comparable to that of ^1-140^αS_acetyl_ (Figure 3B, D); neither ^1-78^αS_acetyl_ or ^1-53^αS_acetyl_ showed significant internalization (Figure 3B, D). Both ^1-78^αS_acetyl_ and ^1-53^αS_acetyl_ show evidence of plasma-membrane localization, but many fewer internal puncta than the longer constructs (Figure 3D). Comparison of the fraction of internalized N1β and αS_acetyl_ following 12 hours of incubation reveals that the majority the fluorescence signal originates from intracellular protein for the three longer αS_acetyl_ constructs, while significantly less of the fluorescence signal comes from intracellular protein for the two shorter αS_acetyl_ constructs (Figure 3E). At the same 12 hour time-point, co-localization analysis of fluorescence from both N1β and αS_acetyl_ on the plasma membrane and in intracellular puncta show in inversion between the three longer and two shorter constructs (Figure 3 Table); for ^1-140^αS_acetyl_, 1-121αS_acetyl_ and ^1-100^αS_acetyl_ a large Pearson’s coefficient for the internalized protein reflects a high degree of co-localization with N1β, while a large Pearson’s coefficient is calculated for plasma-membrane localized ^1-78^αS_acetyl_ and ^1-53^αS_acetyl._ Conversely, the small Pearson’s coefficients calculated for plasma-membrane localized ^1-140^αS_acetyl_, 1-121αS_acetyl_ and ^1-100^αS_acetyl_ and internalized ^1-78^αS_acetyl_ and ^1-53^αS_acetyl_ is due to the relative small AL594 αS_acetyl_ signal as compared to that of N1β (Figure 3E). ^1-90^αS_acetyl_ uptake is comparable to the longer αS_acetyl_ constructs (Supplemental Figure S4), highlighting residues 79-90 as critical for differentiating internalization. Aggregated (pre-formed fibrillar, PFF) ^1-140^αS_acetyl_ (^PFF^αS_acetyl_) is internalized and co-localizes with N1β (Figure 3B). In the absence of N1β expression, none of the αS_acetyl_ constructs were spontaneously internalized by HEK293 cells (Supplemental Figure S5). Spearman’s coefficients calculated for cytoplasmic proteins are consistent with the Pearson’s analysis (Figure 3 Table).

We speculated that the observed differences in internalization between the constructs may be due to reduced binding of the shorter αS_acetyl_ constructs. To test this, we treated HEK-N1β cells with an endocytosis inhibitor (Dynasore hydrate) and took advantage of the fact that even in the absence of Dynasore, relatively little αS_acetyl_ is internalized by HEK-N1β at short time points (<1 hour; Figure 3A). Each of the constructs was incubated with Dynasore treated HEK-N1β cells for 30 minutes to allow for binding with minimal cellular uptake. Confocal imaging was used to quantify the relative signal of membrane-localized αS_acetyl_ normalized to the membrane-localized N1β signal, to account for cell-to-cell variability in N1β expression levels. Surprisingly, we found that all five constructs bind comparably at 200 nM (Figure 3F), indicating that differences in internalization of the various αS_acetyl_ constructs are not due to major differences in their affinities for HEK-N1β cells. The average expression levels of N1β? as measured by the signal intensity of the N1β channel in the binding experiments, is consistent across different imaging wells; thus differences in uptake are not due to significant differences in N1β expression between wells (Supplemental Figure S6). Importantly, our results demonstrate that while binding to N1β is localized to the N-terminal/NAC region of αS_acetyl_, residues 79-90 are required for internalization. In prior work, the C-terminus of αS has been proposed to regulate interactions with membranes, other cellular receptors, as well as with itself (Bussell and Eliezer, 2001; Crowther et al., 1998; Eliezer, 2013; Sevcsik et al., 2011; Zhang et al., 2021). Here, we demonstrate that residues 91-140 also regulate the interaction of αS_acetyl_ with N1β? enhancing N1β-mediated cellular uptake (Figures 2,3).

### Truncation results in inhibited dynamics for N1β/plasma membrane associated αS_acetyl_

In order to gain more insight into the nature of the interactions between αS_acetyl_ and N1β underlying uptake, fluorescence recovery after photobleaching (FRAP) was used to measure the dynamics both of AL594 αS_acetyl_ and N1β−tGFP in the plasma membrane of Dynasore treated HEK-N1β cells (Figure 4A). In the absence of αS_acetyl_, N1β fluorescence recovers ona timescale of ∼3 minutes (Figure 4B). The fluorescence recovery of N1β following the addition of full-length ^1-140^αS_acetyl_, ^1-121^αS_acetyl_ or ^1-100^αS_acetyl_ does not change significantly (Figure 4C); moreover, when the fluorescence of the αS_acetyl_ was measured, the kinetics of the αS_acetyl_ fluorescence recovery tracks with that of N1β recovery for all three constructs.

**Figure 4:**
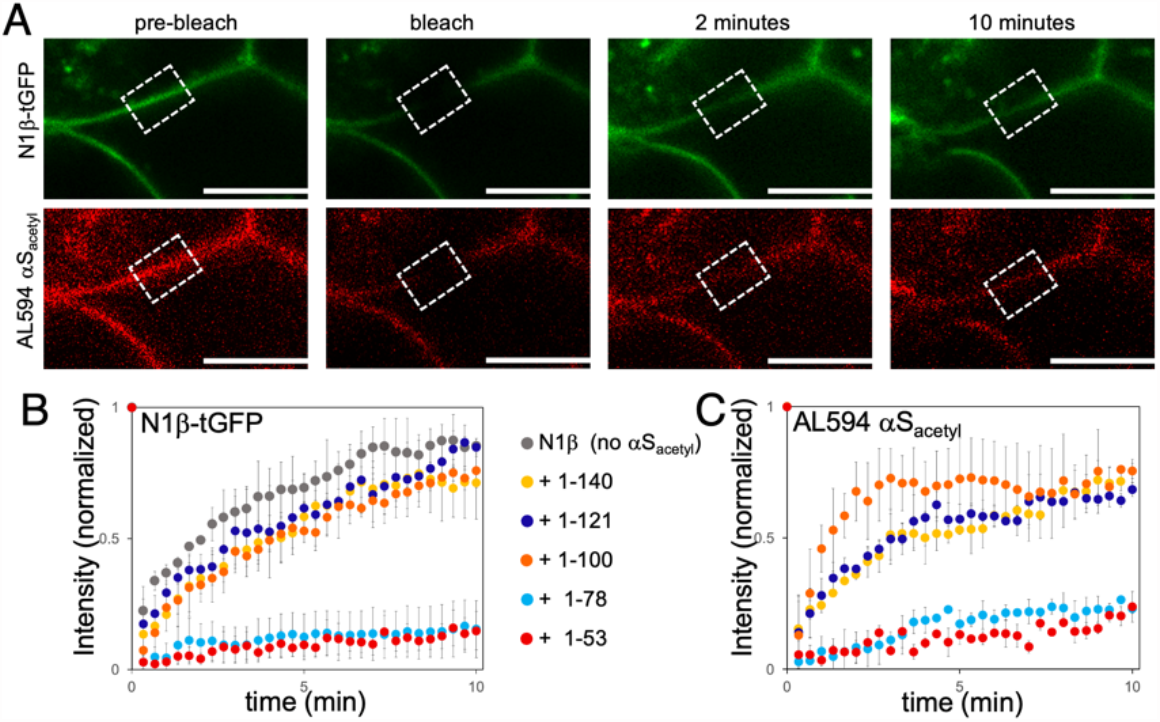
Internalization of αS_acetyl_ is correlated with dynamics on the cell membrane. (A) Representative images of HEK-N1β−tGFP with AL594 ^1-140^αS_acetyl_ bound used for FRAP analysis. (B) Averaged FRAP recovery curves of HEK-N1β (left) and AL594 αS_acetyl_ (right) fluorescence, as labeled. Curves were corrected for steady-state photobleaching and normalized to control fluorescent levels. Two ROIs from three independent experiments were chosen for FRAP. Scale bars = 10 μm.

In striking contrast, the fluorescence recovery of N1β in the presence of either ^1-78^αS_acetyl_ or ^1-53^αS_acetyl_ is dramatically slowed (Figure 4B); as with the other truncated constructs, ^1-78^αS_acetyl_ and ^1-53^αS_acetyl_ show similarly slowed dynamics (Figure 4C) relative to the longer constructs. Overall, the dynamics of diffusion of N1β appears to be correlated with the dynamics of the specific αS_acetyl_ construct; relatively rapid mobility on the cell surface seems essential for the efficient internalization of both proteins. The highly regulated surface mobility of N1β in neurons has been previously reported (Klatt et al., 2021; Neupert et al., 2015); changes in expression levels, membrane localization and neuronal activity alter its diffusion. Moreover, N1β is cleaved by a number of extracellular proteases (Klatt et al., 2021; Trotter et al., 2019). It is possible that binding of some αS_acetyl_ constructs modifies this cleavage, impacting both N1β dynamics and αS_acetyl_-mediated internalization. Here we identify binding of αS_acetyl_ as altering N1β dynamics, and consequently impacting intracellular trafficking of both proteins. Although molecular details of the inhibited diffusion and cellular uptake of the truncated αS_acetyl_ variants remain to be elucidated, our data provides direct evidence that the dynamic diffusion of N1β underlies αS_acetyl_ uptake.

### Truncated αS_acetyl_ alters internalization of ^1-140^αS_acetyl_

Given their apparent ability to inhibit N1β dynamics, we were curious whether binding of the shorter, truncated variants inhibited internalization of full-length αS_acetyl_. HEK-N1β cells were incubated with excess (1 μM total; 800 nM unlabeled protein and 200nM AL594 labeled protein) ^1-78^αS_acetyl_ or ^1-53^αS_acetyl_ for 1 hour to allow for binding; then 200 nM AL647 ^1-140^αS_acetyl_ was added and incubated for an additional 12 hours. ^1-140^αS_acetyl_ was still internalized by HEK-N1β cells, even in the presence of 5-fold excess of ^1-53^αS_acetyl_ (Figure 5A); in fact, under these conditions ^1-140^αS_acetyl_ drives internalization of ^1-53^αS_acetyl_, as seen by co-localization in cellular puncta of both αS_acetyl_ variants, along with N1β (Figure 5A). In contrast, ^1-78^αS_acetyl_ was effective at inhibiting uptake of ^1-140^αS_acetyl_ (Figure 5B). It appears that excess ^1-78^αS_acetyl_ likely blocks the binding site or regions on N1β critical to the internalization of full-length protein. By blocking uptake of ^1-140^αS_acetyl_, ^1-78^αS_acetyl_ also disrupts the uptake of N1β (Figure 5B, C).

**Figure 5:**
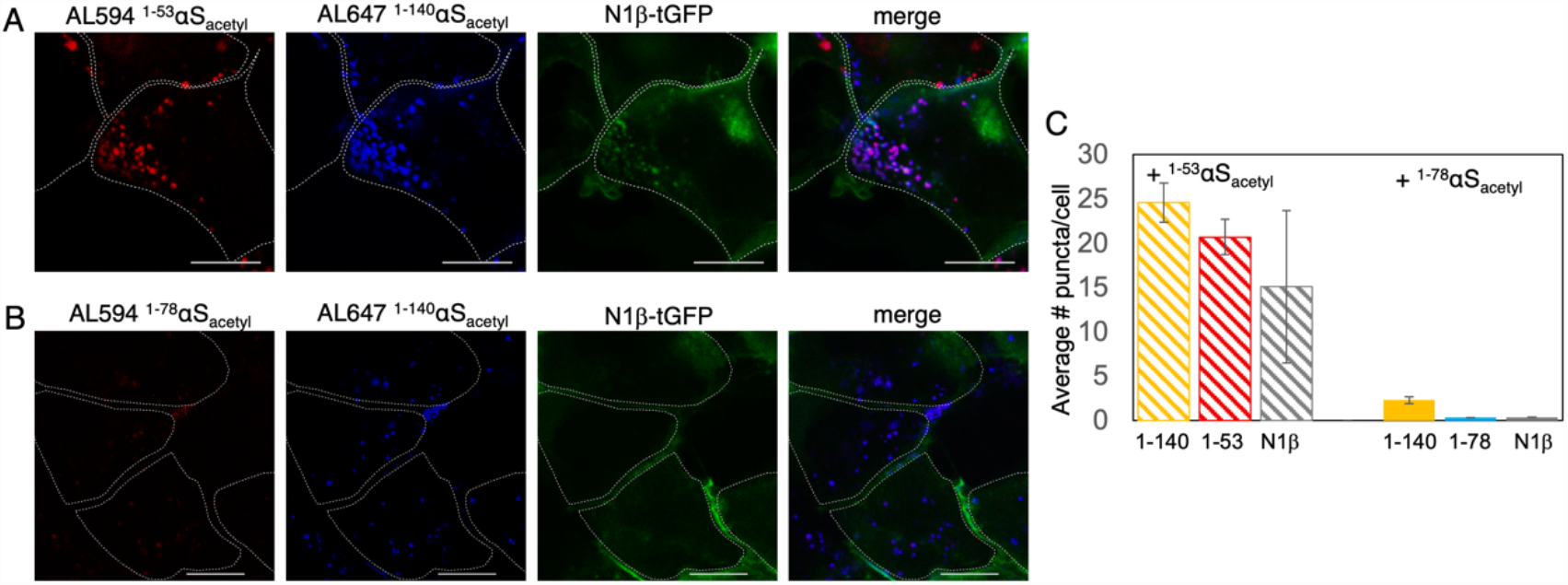
Internalization of^1-140^αS_acetyl_ is diminished by^1-78^αS_acetyl._ Representative images of HEK-N1β cells pre-incubated with AL594^1-53^αS_acetyl_ (A) or AL594^1-78^αS_acetyl_ (B). (C) Quantification of internalization of AL647^1-140^αS_acetyl_ from (A) and (B) by puncta analysis. Analysis based on n = 100 cells, 2 independent experiments. Scale bars = 20 μm.

Through their interactions with binding partners in the post-synaptic membranes, neurexin family of proteins, including N1β? play a critical role in regulating the functional properties of synapses (Südhof, 2017), including formation and modification of synapses (Dean and Dresbach, 2006). Interestingly, a role for αS in synaptic plasticity has also been proposed, based on an early study showing a decrease in αS mRNA transcripts in regions of the zebra finch brain linked to song learning (George et al., 1995). One study found that N1β interacted with αS PFFs (Mao et al., 2016), with a subsequent paper suggesting the binding of αS PFFs could disrupt N1β-mediated synaptic organization (Feller et al., 2023). Our own prior work demonstrated that the interaction between N1β and αS_acetyl_ monomer was chemically selective, dependent upon both the N-terminal acetylation of αS and the N-linked glycan on the extracellular domain of N1β (Birol et al., 2019). Overall, this study contributes molecular insight in the interaction of these two proteins. Our findings highlight the importance of the C-terminus of αS_acetyl_ for mobility of N1β and αS_acetyl_ in the plasma membrane. In neurons, dynamic rearrangements on membranes are critical for synaptic transmission and plasticity. Our work suggests that dynamics of N1β may be regulated by binding of soluble protein ligands, including αS_acetyl_. Interplay between N1β and monomer αS_acetyl_ may even be able to modify synaptic connectivity in neurons. Lastly, we demonstrate that a αS_acetyl_ truncation is capable of blocking binding to N1β by the full-length, physiological αS_acetyl_, as well as inhibiting its subsequent internalization in cells, highlighting a potential approach for future therapeutic interventions.

## Materials and Methods

### αS expression and purification

αS_acetyl_ was expressed in *E. coli* BL21 cells; BL21 stocks containing the N-terminal acetyltransferase B (NatB) plasmid with orthogonal antibiotic resistance were used. The Nat B-BL21 cells were transformed with T7-7 plasmid containing the human αS sequence and grown in the presence of both chloramphenicol (34 μg/mL) and ampicillin (100 μg/mL) to select for both Nat B and αS expression (as described in (Birol et al., 2019) with minor modifications). Briefly, two ammonium sulfate cuts were used (0.116 g/mL and 0.244 g/mL) with αS_acetyl_ precipitating in the second step. The pellet was resolubilized in Buffer A (25 mM Tris pH 8.0, 20 mM NaCl, 1 mM EDTA) with 1 mM PMSF and dialyzed against Buffer A to remove ammonium sulfate.

Dialyzed samples for^1-140^αS_acetyl_ and^1-121^αS_acetyl_ were loaded to an anion exchange column (GE HiTrap Q HP, 5 ml) and eluted with a gradient to 1 M NaCl.^1-140^αS_acetyl_ elutes at approximately 300 mM NaCl. Fractions containing αS were pooled and concentrated using Amicon Ultra concentrators (3,000 Da MWCO). For^1-53^αS_acetyl_,^1-78^αS_acetyl_,^1-90^αS_acetyl_ and^1-100^αS_acetyl_, samples were loaded onto an anion exchange and flow-through was collected and concentrated. For all αS_acetyl_ constructs, concentrated samples were then loaded to a size exclusion column (GE HiLoad 16/600 Superdex75) and eluted at 0.5 ml/minute. Fractions containing αS were again pooled and concentrated, then stored at −80°C. All αS constructs used in this work were checked by matrix-assisted laser desorption/ionization (MALDI) to confirm correct mass and presence of acetylation.

### αS labeling

For site-specific labeling of αS_acetyl_, a cysteine was introduced at either residue 9 or residue 130. For labeling reactions, freshly purified αS_acetyl_ (typically 200–300 μL of approximately 200 μM protein) was incubated with 1 mM DTT for 30 minutes at room temperature to reduce the cysteine. The protein solution was passed over two coupled HiTrap Desalting Columns (GE Life Sciences, Pittsburgh, PA) to remove DTT and buffer exchanged into 20 mM Tris (pH 7.4), 50 mM NaCl, and 6 M guanidine hydrochloride (GdmCl). The protein was incubated overnight at 4°C with stirring with 4× molar excess AL594 or AL647 maleimide (Invitrogen). The labeled protein was concentrated and buffer exchanged into 20 mM Tris pH 7.4, 50 mM NaCl using an Amicon Ultra 3K Concentrator (Millipore, Burlington, MA), with final removal of unreacted dye and remaining GdmCl by passing again over a set of coupled desalting columns equilibrated with 20 mM Tris pH 7.4, 50 mM NaCl.

### Fibril formation

αS_acetyl_ PFFs were prepared as previously described (Volpicelli-Daley et al., 2014). Briefly, 100 μM αS_acetyl_ was mixed with 5 μM AL594 αS_acetyl_ in 20 mM Tris pH 7.4, 100 mM NaCl. To induce aggregation, this solution was incubated at 37°C for 5 days with agitation (1,500 rpm on an IKA MS3 digital orbital shaker) in parafilm-sealed 1.5 mL Eppendorf tubes to ensure minimal solvent evaporation. The aggregation reaction was analyzed by Congo Red absorbance by diluting 10 μl of the aggregation solution in 140 μl 20 μM Congo Red. The mature fibrils were then pelleted by centrifugation (13,200 rpm for 90 minutes at 4°C), and the supernatant was removed. Fibers were resuspended in an equal volume (relative to supernatant) of 20 mM Tris pH 7.4, 100 mM NaCl. Mature fibers were subsequently fragmented on ice using a sonicator (Diagenode UCD-300 bath sonicator) set to high, 30 seconds sonication followed by a delay period of 30 seconds—10 minutes total—to form PFFs.

### Cell culture

SH-SY5Y and HEK293 cells were grown at 37°C under a humidified atmosphere of 5% CO_2_. The SH-SY5Y cells were cultured in Dulbecco’s Modified Eagle’s Medium (DMEM) plus 10% fetal bovine serum, 50 U/ml penicillin, and 50 μg/ml streptomycin. The HEK cells were cultured in DMEM supplemented with 10% FBS, 2 mM L-glutamine, and 100 U/mL each of penicillin and streptomycin. Cells were passaged upon reaching approximately 95% confluence (0.05% Trypsin-EDTA, Life Technologies, Carlsbad, CA), propagated, and/or used in experiments. Cells used in experiments were pelleted and resuspended in fresh media lacking Trypsin-EDTA.

For imaging of SH-SY5Y cells, cells were grown in 8-well ibidi chambers (μ-Slide, 8-well glass bottom, ibidi GmbH, Germany) which were coated with poly-D-lysine the night before.

Chambers were seeded with 20,000–25,000 cells/well and cultured for 48 hours before beginning experiments. For cellular uptake experiments, AL594 labeled αS_acetyl_ was added to cell media to a final concentration of 200 nM. Following incubation (0 to 24 hours), the media was exchanged to remove non-internalized AL594 labeled αS_acetyl_ and cells were allowed to recovery briefly (30 minutes) before acquiring images.

### Transfection of HEK cells

HEK cells were transfected with plasmid encoding the +SS4 isoform of N1β with a C-terminal tGFP tag (Origene) (HEK-N1β) or N1β with an N-terminal tGFP tag (custom plasmid from Origene) by Lipofectamine 3000, following the manufacturer’s directions. The HEK-N1β cells were grown to 70% confluency on 8-well ibidi chambers before transfection. Transfections were performed with 100 ng final concentration of N1β-tGFP or tGFP-N1β plasmid per 250 μL media per well. Transfection media was removed from cells 3 hours later and fresh media was added. At 48 hours following transfection, the media was exchanged with fresh media prior to the addition of αS. Cells were incubated with AL594 or AL647 αS_acetyl_ monomer (final concentration 200 nM) or PFFs (final concentration 200 nM monomer units, 1:20 labeled:unlabeled) for the indicated time points. Cells were then washed with fresh media prior to imaging.

For binding and FRAP experiments, cells were pretreated with the dynamin-dependent endocytosis blocker, Dynasore (Sigma-Aldrich, D7693), at 80 μM, 30 minutes prior to the addition of AL594 αS_acetyl_.

### Lysotracker

For colocalization of AL594 αS_acetyl_ and N1β-tGFP with lysosomes, cells were treated with 75 nM Lysotracker Deep Red (Life Technologies, Carlsbad, CA) for 1 hour prior to imaging.

### Internalization blocking experiments

HEK-N1β cells were incubated with 1 μM (800nM dark and 200nM AL594 labeled)^1-53^αS_acetyl_ or^1-78^αS_acetyl_ for 1 hour, followed by addition of 200 nM AL647^1-140^αS_acetyl._ The cells were incubated for another 12 hours and then washed with media before imaging.

### Cell imaging and analysis

All cell imaging was carried out by confocal fluorescence microscopy using an Olympus FV3000 scanning system configured on a IX83 inverted microscope platform with a 60× Plan-Apo/1.1-NA water-immersion objective with DIC capability (Olympus, Tokyo, Japan). For all experiments, the gain setting for the channels was kept constant from sample to sample: for detection of N1β-tGFP, blue channel excitation 488 nm, emission BP 500–540 nm, for detection of AL594 αS_acetyl_, green channel excitation 561 nm, emission BP 570–620 nm and for detection of AL647 αS_acetyl_ or LysoTracker Deep Red, red channel excitation 640 nm, emission BP 660-720 nm.

Most image acquisition and processing were performed with the software accompanying the FV3000 microscope and Image J software(Schneider et al., 2012). For SH-SY5Y cells, internalized αS_acetyl_ was quantified either by analysis of the punctate structures in the cells or by the total cellular fluorescence. For total cellular fluorescence, the integrated fluorescence intensity of the cells is reported. Cellular puncta were analyzed using the Image J particle analysis plug-in. This algorithm detects puncta through a user-defined threshold and counts the number of puncta that meet or exceed the threshold. The threshold was initially defined by manual identification and averaging of a subset of puncta. Colocalization with N1β was computed by obtaining a Pearson or Spearman coefficient using the ImageJ plugin for colocalization (Coloc_2). The total intensity of AL594 αS_acetyl_ and N1β-tGFP on the cell membrane and inside the cell was quantified by masking the cell membrane and then defining the inside of the masked area as the cell interior. The ImageJ ‘measure’ command was used to get the average intensity integrated over each masked area (cell membrane) and the cell interior separately. Colocalization analysis was carried out on 5-10 frames of cell images, each frame containing at least 5 cells for a total of 25-50 cells.

For some of the experiments, imaging was carried out on a PicoQuant MicroTime 200 time-resolved fluorescence system based on an inverted Olympus IX73 microscope (Olympus, Tokyo, Japan). A 60X Plan-Apo/1.4-NA water-immersion objective, 482 nm and 560 nm excitation lasers and a frame size of 512 × 512 pixels was used. Images acquired with this instrument were in lifetime mode but were integrated to obtain intensity-based images comparable to typical confocal images. Fluorescence intensities were analyzed via the lifetime mode using SymPhoTime 64 (PicoQuant, Berlin, Germany). The intensity of images was then adjusted on ImageJ analysis program.

### FRAP

FRAP was performed on the same Olympus confocal microscope described above. A 128 ×128 pixel image was captured at 0.188 s intervals using the 488 nm and 561 nm laser lines at 1% power and the pinhole set to 105 μm. Following 20 prebleach frames, a 50×50 pixel region was bleached at full power for 0.94 seconds (one scan iteration). Images were captured until no further recovery was evident. Image stacks were loaded, and both the rough center of the bleach area and an appropriate unbleached reference area were selected manually. Mean fluorescence was monitored within the bleached region as well as within the manually specified reference region to control for photobleaching. Individual recovery curves were normalized to the reference curves, and then to prebleach values.

## Supporting information

SuppInfo

## Abbreviations

N1β: neurexin 1β
αS: α-Synuclein
AL594: Alexa Fluor 594 C_5_ maleimide
AL647: Alexa Fluor 647 C_5_ maleimide
acetyl: N-terminally acetylated

## Acknowledgements

We thank T. Baumgart for use of his confocal microscope and V. M.-Y. Lee for use of her sonicator. This work was supported by the University of Pennsylvania and the National Institutes of Health (R01 NS120625 to E.R.) and MDC-Berlin (I.I.D.M.).

