## Supplementary material for "The C-terminus of α-Synuclein regulates its dynamic cellular internalization by Neurexin 1β": SuppInfo

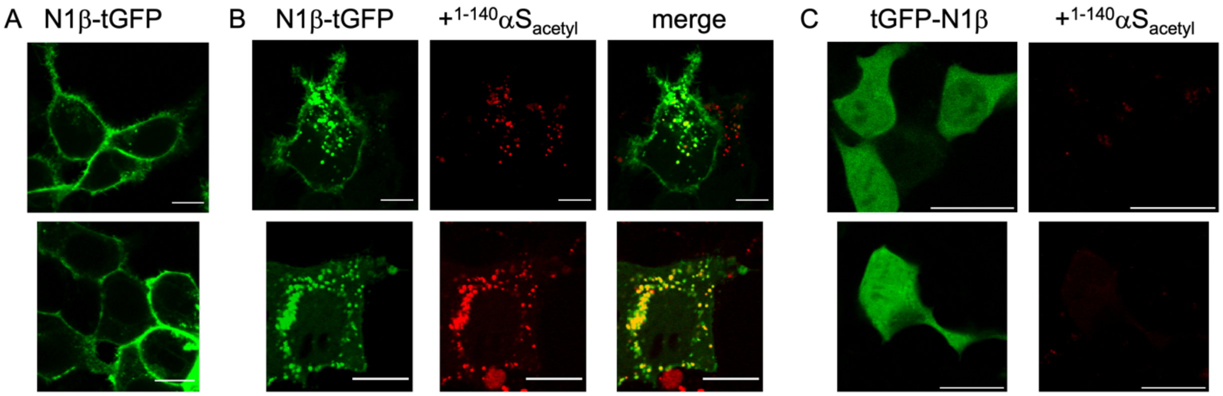


**Figure S1: N1β–tGFP localizes to the plasma membrane** (A) Following 48 hours transfection, the majority (~75 %; see quantification in Figure S2) of N1β–tGFP localizes to the plasma membrane of HEK293 cells; this is consistent with a number of other studies using N1β with a C-terminal fluorescent protein tag in functional studies of N1β (Graf et al., 2006; Graf et al., 2004; Khalaj et al., 2020; Siddiqui et al., 2010; Sterky et al., 2017). (B) Following 12 hours of incubation with AL594 $^{1\text{-}140}\alpha S_{acetyl}$, some N1β–tGFP can still be observed at the plasma membrane, but it can also be observed co-localized with AL594 $^{1\text{-}140}\alpha S_{acetyl}$ in intracellular puncta. (C) tGFP-N1β is soluble and localized throughout the cell body, due to blocking of the signal sequence. Following 12 hours of incubation with AL594 $^{1\text{-}140}\alpha S_{acetyl,}$ HEK293 cells expressing tGFP-N1β do not show any detectable internalization of AL594 $^{1\text{-}140}\alpha S_{acetyl}$.

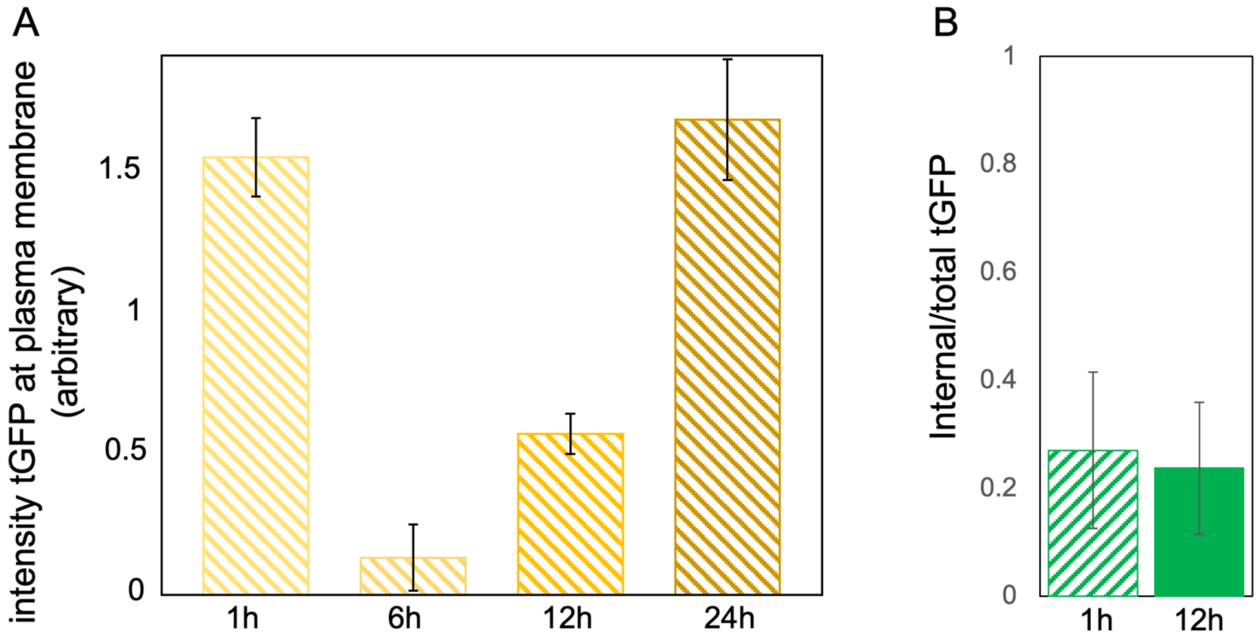


**Figure S2: N1β -tGFP is recycled to the plasma membrane following internalization.** (A) Quantification of N1b-tGFP on the cell membrane of HEK cells following 1h, 6h, 12h, and 24h incubation with AL594 $^{1-140}\alpha S_{acetyl}$. The total intensity of N1β-tGFP on the cell surface is reported. This plot only quantifies the intensity at the plasma membrane, not the total amount of N1β present. Some fraction of N1β remains internalized and co-localized with $^{1-140}\alpha S_{acetyl}$ at 24 hours, as can be seen in Figure 3A. Analysis for is based on n = 100 cells, 3 independent experiments. (B) Quantification of N1β-tGFP signal from cell cytoplasm. In the absence of $\alpha S_{acetyl}$ incubation, the fraction of cytoplasmic N1β-tGFP signal remains constant at ~25%. The 12 h time point is also plotted in Figure 3E. Colocalization analysis based on n = 25 – 50 cells, 3 independent experiments.

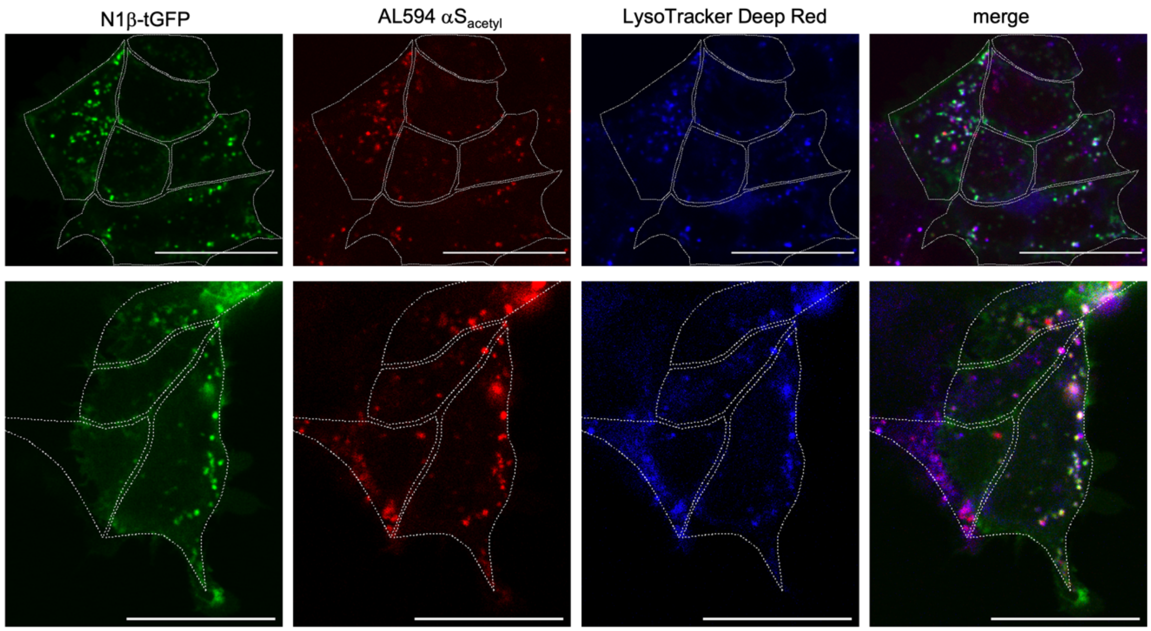


**Figure S3: Internalized N1β–tGFP and AL594 $^{1\text{-}140}$αS$_{acetyl}$ co-localize with LysoTracker Deep Red.** Images are taken in HEK- N1β cells, following 12 hours of incubation with AL594 $^{1\text{-}140}$αS$_{acetyl}$ and 1 hour incubation with LysoTracker Deep Red.

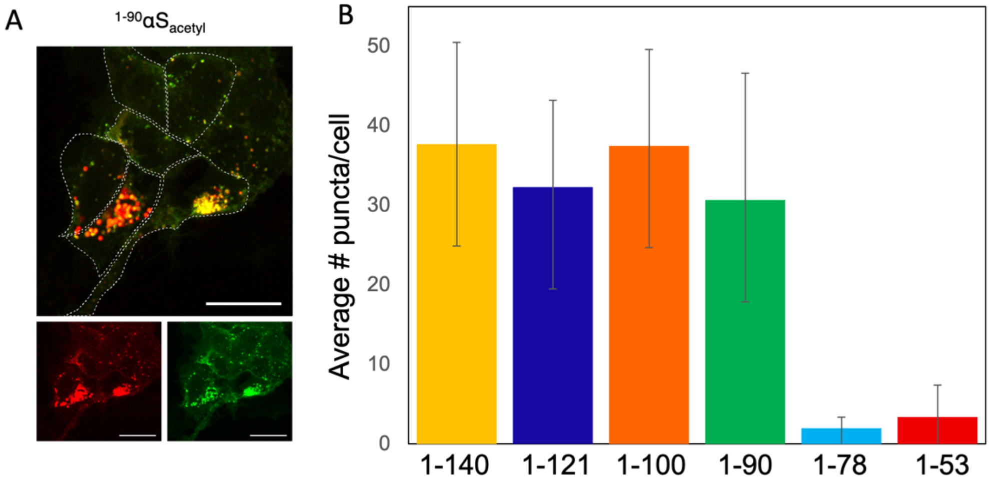


**Figure S4: Internalization of $^{1-90}\alpha S_{acetyl}$ is similar to $^{1-100}\alpha S_{acetyl}$.** (A) Representative image of HEK-N1β-tGFP cells following 12h incubation with AL594 $^{1-90}\alpha S_{acetyl}$. The αS (red) and N1β (green) channels are shown separately below a larger image of their merge; cells are outlined in white dashed lines. (B) Quantification of internalization of AL594 $^{1-90}\alpha S_{acetyl}$ by HEK-N1β cells following 12 h incubation quantified by puncta analysis. This is a replicate of Figure 3D, with the $^{1-90}\alpha S_{acetyl}$ added to allow for comparison. Analysis for internalization is based on n = 100 cells, 3 independent experiments. Scale bars = 20 μm.

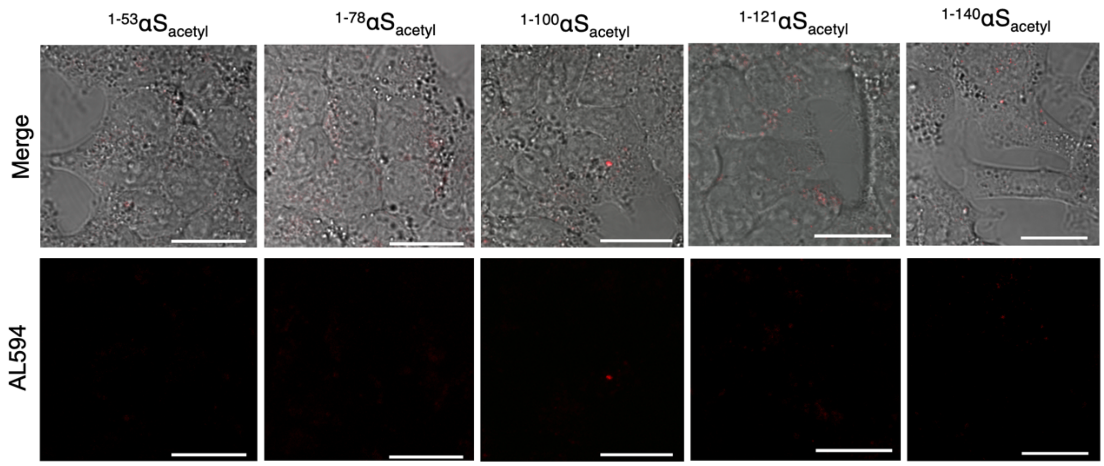


**Figure S5: HEK cells do not spontaneously internalize any $\alpha S_{acetyl}$ variants in the absence of N1β.** Representative images of HEK cells following 12h incubation with AL594 $\alpha S_{acetyl}$ variants, as noted. The merge of fluorescence and DIC and αS (red) channels are shown. Scale bars = 20 μm.

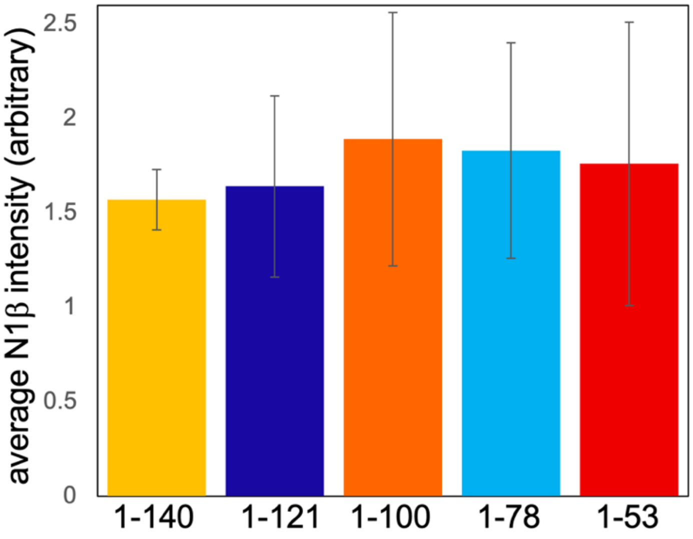


**Figure S6: N1β transfection efficiencies in HEK cells are consistent across wells.** Quantification of N1β-tGFP expression in HEK cells following transfection was quantified by measuring the average intensity of N1β-tGFP on the same samples used for the binding experiments shown in Figure 3. The majority of N1β-tGFP is localized to the plasma membrane. Analysis is based on n = 100 cells, 3 independent experiments.
